# The TFIID subunit Taf4 is required for pancreatic beta cell function and identity

**DOI:** 10.1101/2020.05.23.111898

**Authors:** Thomas Kleiber, Guillaume Davidson, Gabrielle Mengus, Igor Martianov, Irwin Davidson

## Abstract

We selectively inactivated the Taf4 subunit of general transcription factor TFIID in adult murine pancreatic beta cells (BCs). Taf4 inactivation rapidly diminishes expression of critical genes involved in BC function leading to increased glycaemia, lowered plasma insulin levels, defective glucose-stimulated insulin secretion and in the longer term reduced BC mass through apoptosis of a subpopulation of BCs. Nevertheless, glycaemia and blood insulin levels are stabilised after 11 weeks with mutant animals showing long term survival. Bulk RNA-seq and ATAC-seq together with single cell RNA-seq on isolated Langerhans islets show that Taf4 loss leads to a remodelling of chromatin accessibility and gene expression not only in targeted BCs, but also alpha and delta cells. One week after Taf4-loss, cells with mixed BC, alpha and/or delta cell identities were observed as well as a BC population trans-differentiating into alpha-like cells. Computational analysis defines how known critical BC and alpha cell determinants may act in combination with additional transcription factors and the NuRF chromatin remodelling complex to promote BC trans-differentiation.

## Introduction

The endocrine compartment of the pancreas organized in the Langerhans islets comprises multiple hormone-secreting cell types such as insulin-secreting beta cells (BCs), glucagon-secreting alpha cells (ACs), somatostatin-secreting delta cells (DCs) as well as the rarer pancreatic polypeptide-secreting PP cells and ghrelin-secreting epsilon cells. These cells cooperate to regulate glucose homeostasis in the organism.

Plasticity and trans-differentiation between these endocrine cell populations has been described in diabetes, in genetically modified mice and after various pathological or chemical treatments ^1, 2, 3, 4, 5, 6^. ACs appear to have a highly plastic epigenetic state with many loci showing bivalent chromatin modifications ^7^ and ACs can be converted to beta-like insulin secreting cells by overexpression of BC-determining factors ^8^ or loss of the Arx and Dnmt1 factors ^9^. In type 2 diabetes (T2D), hyperglycaemia seems to induce BC dedifferentiation, involving reversion to a Ngn3-expressing precursor state, with diminished expression of BC function genes and key BC transcription factors along with expression of BC forbidden genes ^1, 10, 11, 12, 13^. However, with some exceptions ^9, 14, 15^, the transcriptional mechanisms underlying changes in cell identity are not fully understood.

Proper regulation of gene expression by requires a complex and dynamic dialogue between transcription factors binding promoters and enhancers, chromatin remodelling and modification complexes and the basal transcription machinery ^16, 17, 18, 19, 20^. The multi-subunit basal transcription complex TFIID plays a critical role in this communication. TFIID comprises the TATA-box binding protein (TBP) and 13-14 TBP-associated factors (TAFs) and plays a unique role in pre-initiation complex (PIC) formation ^21, 22^. Recent electron microscopy studies showed that TFIID is organised in three lobes A-C that undergo topological reorganisation during the initial steps of PIC formation as it binds promoter DNA along with TFIIA and TFIIB ^22, 23, 24^. The histone-like TAF4-TAF12 heterodimer is crucial for the structural integrity of lobes A and B ^25, 26^. In lobe B, a region of the conserved TAF4 C-terminal domain makes contacts with promoter DNA and both TAF4 and TAF12 contact the TFIIA-TBP module suggesting that they promote TBP DNA binding and fix the distance between the TBP binding site and the transcription start site.

Genetic knockout of Taf4 in mouse, as is the case with several other TAFs, leads to highly specific effects on PIC formation and gene expression. This may in part be explained by redundancy with its paralog TAF4B that also heterodimerizes with TAF12 and integrates into TFIID, thus maintaining TFIID integrity in absence of TAF4 ^27, 28^. We used somatic inactivation to address the function of murine Taf4 in embryonic and adult murine epidermis or neonatal hepatocytes ^26, 27, 29, 30, 31^ and used germline inactivation to address its role during embryogenesis ^28^. In each biological context, Taf4 regulated specific gene expression programs and functions.

To evaluate the role of Taf4 in BCs, we used Insulin-Cre-ER^T2^ transgenics to inactivate Taf4 in murine adult BCs. We show that Taf4 regulates an extensive gene expression program including critical components of the insulin signalling pathway. Taf4 loss also impacts BC identity and computational analysis of single cell transcriptomic data defined how known critical BC and AC determinants may act in combination with additional transcription factors and the NuRF chromatin remodelling complex to promote BC trans-differentiation into alpha-like glucagon-expressing cells.

## Material and Methods

### Genetically engineered mice

Mice with the floxed Taf4 alleles as previously described ^27^ were crossed with previously described ^32^ *Ins*-Cre-ER^T2^ transgenics and Taf4 inactivated by subcutaneous injection 1 time per day for 3 days with 5 ul of Tamoxifen diluted in oil (final concentration 10mg/ml). Blood glucose and insulin levels were measured over a period of 20 weeks with one measurement per week on the same day at the same time. Glycaemia was measured using a glucometer and insulin levels using the Milliplex Map Kit (Mouse Metabolic Magnetic Bead Panel. MMHMAG-44K, Millipore) Milliplex Map Kit. To assess glucose tolerance, mice were fasted for 12 hours before the first glucose measure and then injection of 100 μl of glucose solution (t= 0, 2g of glucose/ kg mouse weight). Insulin levels were monitored in blood samples taken over a 30-minute period starting 5 minutes before the glucose injection. All animal experiments were performed in accordance with European and national guidelines and policies (2010/63/UE directive and French decree 2013-118) and with approval of the National Ethics Committee.

### Immunostaining and counting of islets

Pancreas were isolated and fixed overnight in 4% paraformaldehyde, washed with PBS, dehydrated, paraffin embedded, and sectioned at 5 μm. For antigen retrieval, the sections were boiled for 20 min in 10 mM of sodium citrate buffer. Sections were permeabilized with 3 × 5 min 0.1% Triton in PBS, blocked for 1 hr in 5% Neutral Goat Serum (NGS) in PBS, and incubated overnight in 5% NGS with primary antibodies: mouse anti-Glucagon [K79bB10] (ab10988), Guinea Pig anti-Insulin (Dako IR00261-2) and mouse anti-TAF4 (TAF II p135) (Santa Cruz (sc-136093). Sections were washed 3 × 5 min 0.1% Triton in PBS, and incubated with secondary antibodies: Goat anti-Guinea pig IgG H&L Alexa Fluor® 647 (ab150187) and Goat anti-Mouse IgG H&L Alexa Fluor® 488 (ab150117) for 2 hr. Sections were subsequently incubated with 1/2000 Hoechst nuclear stain for 10 min, washed 3 × 5 min in PBS, dried and mounted with Vectashild. Islet counting was performed on hematoxyline/eosin (HE) labeled sections of the pancreas. The entire pancreas was sectioned (5 μm slice) and one slice every 300 μm was scanned and analyzed with NanoZoomer Digital Pathology view annotation software to count and measure the islets.

### Isolation of Langerhans islets

Anesthetized mice (40 μL ketamine (100mg/mL) + 24 μL xylazine (20mg/mL) + 160 μL NaCl intraperitoneal injection per mouse) were injected in the pancreas with 4 ml of collagenase solution (7 ml HBSS, 70 μL HEPES (10 μM) (Gibco), 16,5μL DNAse (10mg/mL) (sigma DN5), 14 mg collagenase (2 mg/mL) (sigma Type V), and for isolation of RNA 2,3 μL of RNAsine (40U/μL). The pancreas was dissected and placed a tube containing 2 ml collagenase solution and incubated at 37 minutes at 38°C before stopping with 30mL Quenching Buffer (QB; 122 mL HBSS + 2,9 mL HEPES (25 μM) + 0,6 g BSA (0,5%) and 2 cycles of centrifugation at 1200 rpm for 2 min at 4°C. The pellet was suspended in 10 ml of Histopaque 1100 (5 ml histopaque 1077; Sigma 10771 and 6 ml histopaque 1119; Sigma 11191), centrifuged for 20 minutes at 1500 rpm at 4°C and the pellet washed 2 times with QB. The supernatant was discarded and pelleted islets transferred to a bacterial culture dish to be collected by pipetting. Purified islets were then cultured for 24 hours in RPMI1640 (HEPES 25mM) + final 2 mM L-glutamine (Gibco 11875-093) + 10% FCS 95 + 1% Penicillin/Streptomycin. Intracellular ATP concentration was measured with the Luminescent ATP Detection Assay Kit (ab113849) on isolated islets that were further digested to obtain a single cell population and FACS sorted to normalize the numbers of cells used.

### Bulk RNA-seq from isolated islets

RNA was extracted from isolated islets from 2 mice per genotype with Nucleospin RNA plus XS (Macherey-Nagel). Complementary DNA was generated and linearly amplified from 3 ng total RNA using the Ovation RNA-seq V2 system (NuGEN technologies Inc., Leek, The Netherlands), according to the manufacturer’s instructions. The amplified cDNA was then purified using Agencourt AMPure XP beads (Beckman-Coulter, Villepinte, France) in a 1.8:1 bead to sample ratio and fragmented by sonication using a Covaris E220 instrument (with duty cycle: 10%, maximum incident power: 175 watts and cycles/burst: 200 for 120 seconds). The RNA-seq libraries were generated from 100 ng fragmented cDNA using the Ovation Ultralow v2 library system (NuGEN technologies Inc., Leek, The Netherlands) according to the manufacturer’s instructions, with only 6 PCR cycles for library amplification. The final libraries were verified for quality and quantified using capillary electrophoresis before sequencing on an Illumina Hi-Seq4000. Reads were preprocessed using cutadapt version 1.10 in order to remove adapter and low-quality sequences (Phred quality score below 20). After this preprocessing, reads shorter than 40 bases were discarded. Reads were mapped to rRNA sequences using bowtie version 2.2.8, and reads mapping to rRNA sequences were removed. Reads were mapped onto the mm9 assembly of Mus musculus genome using STAR version 2.5.3a. Gene expression quantification was performed from uniquely aligned reads using htseq-count version 0.6.1p1, with annotations from Ensembl version 67 and “union” mode. Only non-ambiguously assigned reads were retained. Read counts were normalized across samples with the median-of-ratios method ^33^. Comparisons of interest were performed using the Wald test for differential expression ^34^ and implemented in the Bioconductor package DESeq2 version 1.16.1. Genes with high Cook’s distance were filtered out and independent filtering based on the mean of normalized counts was performed. P-values were adjusted for multiple testing using the Benjamini and Hochberg method [**3**]. Heatmaps were generated with R-package pheatmap v1.0.12. Deregulated genes were defined as genes with log2(fold change) >1 or <−1 and adjusted p-value <0.05. Gene ontology analyses was performed using GSEA (https://www.gsea-msigdb.org/gsea/index.jsp) or David (https://david-d.ncifcrf.gov/).

### ATAC-seq from isolated islets

ATAC-seq was performed from 20 000 cells from isolated islets. Sequenced reads were mapped to the mouse genome assembly mm9 using Bowtie ^35^ with the following arguments: “-m 1 --strata --best -y -S -l 40 -p 2”.

After sequencing, peak detection was performed using the MACS software ^36^ v2.1.1.20160309. Peaks were annotated with Homer(http://homer.salk.edu/homer/ngs/annotation.html) using the GTF from ENSEMBL v67. Peak intersections were computed using Bedtools ^37^ Global Clustering was done using seqMINER ^38^. In silico footprinting signatures were calculated using TOBIAS ^39^ (v0.5.1; https://github.com/loosolab/TOBIAS/), differential footprinting scores were plotted with R-package ggplot2 (https://ggplot2.tidyverse.org.).

### Single cell RNA-seq from isolated islets

After islet isolation, further digestion was performed with Accutase (A6964, Sigma) at 27 °C for 5 minutes and the cells were suspended in culture medium. Cells were then FACS sorted to select only live cells. 300 islets from a male WT mouse and 300 islets pooled from 2 male mutant mice 1 week or 5 weeks after Taf4 inactivation were used for scRNA-seq. Cell capture was performed using 10X Genomics Chromium Analyzer. After sequencing, raw reads were processed using CellRanger (v 3.1) to align on the mm10 mouse genome, remove unexpressed genes and quantify barcodes and UMIs. Data were then analyzed in R (v3.6.3). For the WT sample only, potential doublets were removed by manually filtering out cells expressing high levels of Ins1+Gcg, Ins1+Sst or Gcg+Sst and then by running R-package DoubletFinder ^40^. The WT sample was down-scaled to 12000 cells before aggregation with Taf4^b−/−^ 5-week sample and analyzed using Seurat v3.1.4 ^41^ following the guided clustering tutorial vignette. Only cells with feature count ranging from 200 to 1500 and with percentage of mitochondrial reads <15% were kept for the analysis. Counts are normalized with the “LogNormalize” method and data are scaled to remove unwanted sources of variation. Clustering was done on most variable features using 12 PCs and a resolution of 0.8. Regulome analyses were performed using the SCENIC v1.1.2.2 package.

## Results

### Inactivation of Taf4 in adult pancreatic beta cells leads to hyperglycaemia

To inactivate Taf4 selectively in adult pancreatic BCs, mice with the previously described floxed Taf4 alleles ^26, 27^ were crossed with an *Ins1*::Cre-ER^T2^ transgenic driver ^32^ to generate *Ins1*::Cre-ER^T2^::*Taf4*^lox/lox^, *Taf4*^lox/+^ or *Taf4*^+/+^ animals. At 11-13 weeks of age, adult animals were injected with Tamoxifen (Tam) on 3 consecutive days to generate *Taf4*^b−/−^ mice lacking Taf4 in BCs and control animals from the other genotypes that maintained Taf4 expression. Immunostaining of Langerhans islets revealed loss of Taf4 expression already 1 week after injection in *Taf4*^b−/−^ animals, whereas it was clearly visible in Tam injected *Ins1*::Cre-ER^T2^::*Taf4*^+/+^ controls (Fig. 1A). We noted also the absence of Taf4 in the nuclei of the acinar cells surrounding the islets in both the control and *Taf4*^b−/−^ mutant animals suggesting that these cells normally did not express Taf4 (Fig. 1A and B). No Taf4 expression was detected in Langerhans islets of the *Taf4*^b−/−^ animals 3 and 10 weeks after injection, but strikingly its expression was detected in a sub-population of BCs after 34 weeks (Fig. 1A and B). After 55 weeks, islets were still composed of a mix of Taf4 expressing and non-expressing BCs, showing that even after this prolonged period the Taf4 null population was not completely replaced by Taf4 expressing cells. In each case, cells that lacked or regained Taf4 expression showed cytoplasmic insulin labelling indicating they were insulin producing BCs. This observation suggested that over time the BC population was partially reconstituted from cells that either had not undergone recombination of the Taf4 alleles, from a precursor cell population or trans-differentiation of other Langerhans islet cell populations into BCs. These data showed that Taf4 can be efficiently inactivated in BCs and that Taf4 mutant BCs were replaced only partially over the long term by a novel population expressing Taf4.

**Figure 1.**
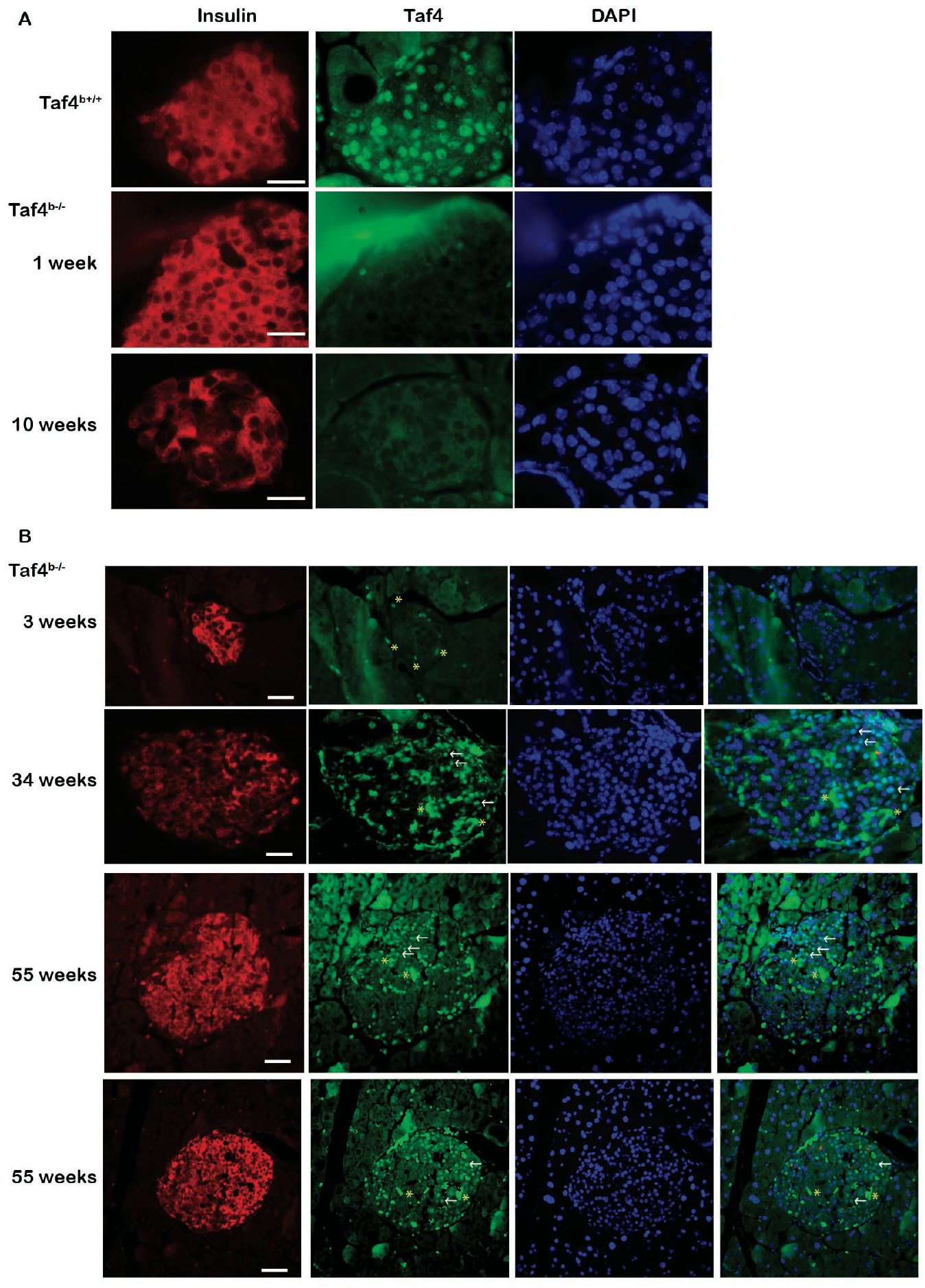
Inactivation of Taf4 in BCs. **A**. Immunostaining of isolated Langerhans islets from mice with the indicated genotypes for Taf4 or insulin as indicated. The number of weeks after Tam injection are indicated. **B**. Immunostaining of isolated Langerhans islets as above but with the addition of the Taf4-DAPI merge. Representative Taf4-positive nuclei at 34 and 55 weeks are indicated by arrows, not to be confounded with the stronger non-specific staining indicated by *. Scale bar = 100 μM.

We examined evolution of different physiological parameters over a period of 20 weeks following Taf4 inactivation. Measurement of blood glucose levels in ad libitum-fed mice indicated, that while no change was seen 1 week after Taf4 loss, a potent increase in glycaemia was observed after 3 weeks (Fig. 2A). Hyperglycaemia persisted over the 20 weeks, although levels were mildly but reproducibly reduced after 10 weeks compared to what was observed at 3 and 6 weeks. In agreement with this, circulating plasma insulin levels were also reduced between weeks 1-10, but recovered somewhat at later times (Fig. 2B). Measurement of ATP levels, a regulator of insulin secretion, in wild-type and mutant islets showed a strong decrease 3 weeks after Tam injection in the *Taf4*^b−/−^ animals, but a progressive recovery to almost normal levels between 11 and 24 weeks (Fig. 2C). Loss of Taf4 in BCs therefore resulted in increased glycaemia and lowered plasma insulin levels, that were partially, but not fully, restored after 10 weeks. As a result, the *Taf4*^b−/−^ animals did not show a total loss of BC function and no premature death with lifespans comparable to their Taf4 expressing littermates.

**Figure 2.**
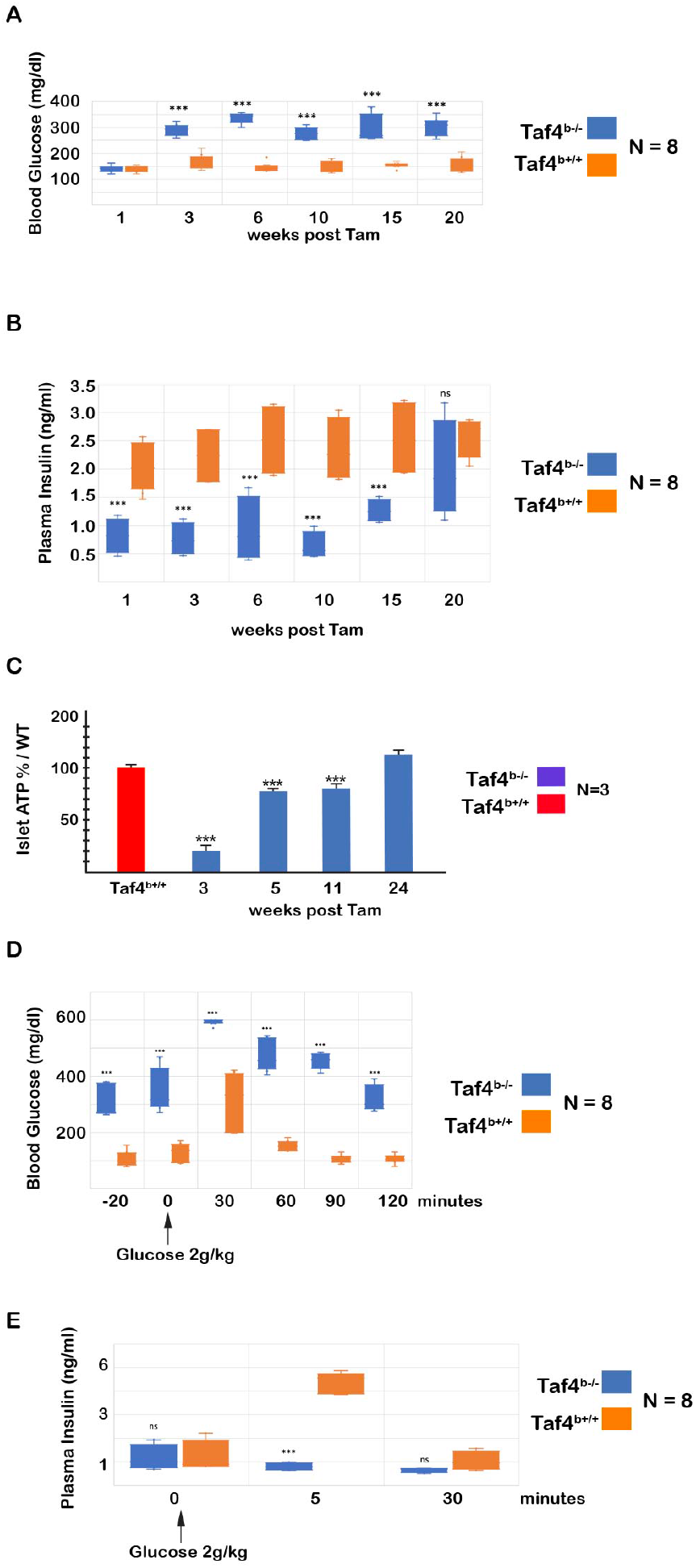
Physiological parameters of *Taf4*^*b−/−*^ animals. **A-B**. Blood glucose and insulin levels of *Taf4*^+/+^ and *Taf4*^*b−/−*^ animals at the indicated number of weeks following Tam injection. **C.** ATP content of isolated islets from control animals or from *Taf4*^*b−/−*^ animals at the indicated number of weeks following Tam injection. **D**. Blood glucose levels at the indicated times following injection of glucose in control and *Taf4*^*b−/−*^ animals. Mice at 14 weeks were injected with Tam and glucose administered 3 weeks following Tam treatment. **E.** Plasma insulin levels from the same protocol. Student T-test *** p<0.001 from N=8.

### Defective glucose-stimulated insulin secretion upon Taf4 inactivation

To assess whether Taf4 loss lead to defective glucose-stimulated insulin secretion (GSIS) and glucose tolerance, *Taf4*^b−/−^ animals and their Taf4-expressing littermates 3 weeks after Tam injection were fasted overnight and then injected with glucose and the resulting glycaemia and plamsa insulin levels were monitored. Before injection, the *Taf4*^b−/−^ animals showed the increased glycaemia seen after 3 weeks that was further increased 30 minutes after glucose administration (Fig. 2E). Compared to their Taf4-expressing littermates, glycaemia in the *Taf4*^b−/−^ animals increased to higher levels that persisted over 90 minutes before returning to basal values by 120 minutes. In contrast, control animals showed a lower increase in glycaemia that returned to basal values after 90 minutes. Similarly, glucose administration in *Taf4*^b−/−^ animals failed to increase plasma insulin levels as seen in control animals (Fig. 2F). Taf4 inactivation therefore led to diminished GSIS and altered glucose tolerance.

### Taf4 inactivation leads to defective insulin maturation and apoptosis of a subset of beta cells

We examined BC fate following loss of Taf4. Histological examination of pancreatic sections allowed quantification of the number and size of Langerhans islets. At 3 weeks, the number of islets was comparable in *Taf4*^b−/−^ and control animals, however, by 11 weeks their numbers were significantly lower in the knockouts (Fig. 3A). We additionally crossed the *Ins1*::CreER^T2^::*Taf4*^lox/lox^ animals with mice bearing a Lox-Stop-Lox cassette driving a GFP reporter in the *Rosa26* locus such that Tam administration activated GFP expression in BCs that were subsequently isolated from islets by FACS. After 11 weeks, a 7-fold reduction in BCs was observed following Taf4 loss (Fig. 3A). Consistent with the loss of BCs, at 3 weeks, and more strikingly at 11 weeks, the % of islets of smaller size were increased in the *Taf4*^b−/−^ animals (Fig. 3B and C). Staining of islets by TUNEL at 3 weeks in both *Taf4*^b−/−^ and control islets showed few apoptotic cells at the periphery and only rare apoptotic BCs within the islet (Fig. 3D). After 11 weeks, numerous apoptotic BCs were seen within the islets. Quantification indicated that by 3 or 11 weeks after Tam injection 100% of the *Taf4*^b−/−^ islets comprised apoptotic BCs within the islet, but only 30% of the control animals (Fig. 3E). Taf4 loss therefore led to a progressive loss of BCs through apoptosis that began around 3 weeks, but that was strongly amplified by 11 weeks.

**Figure 3.**
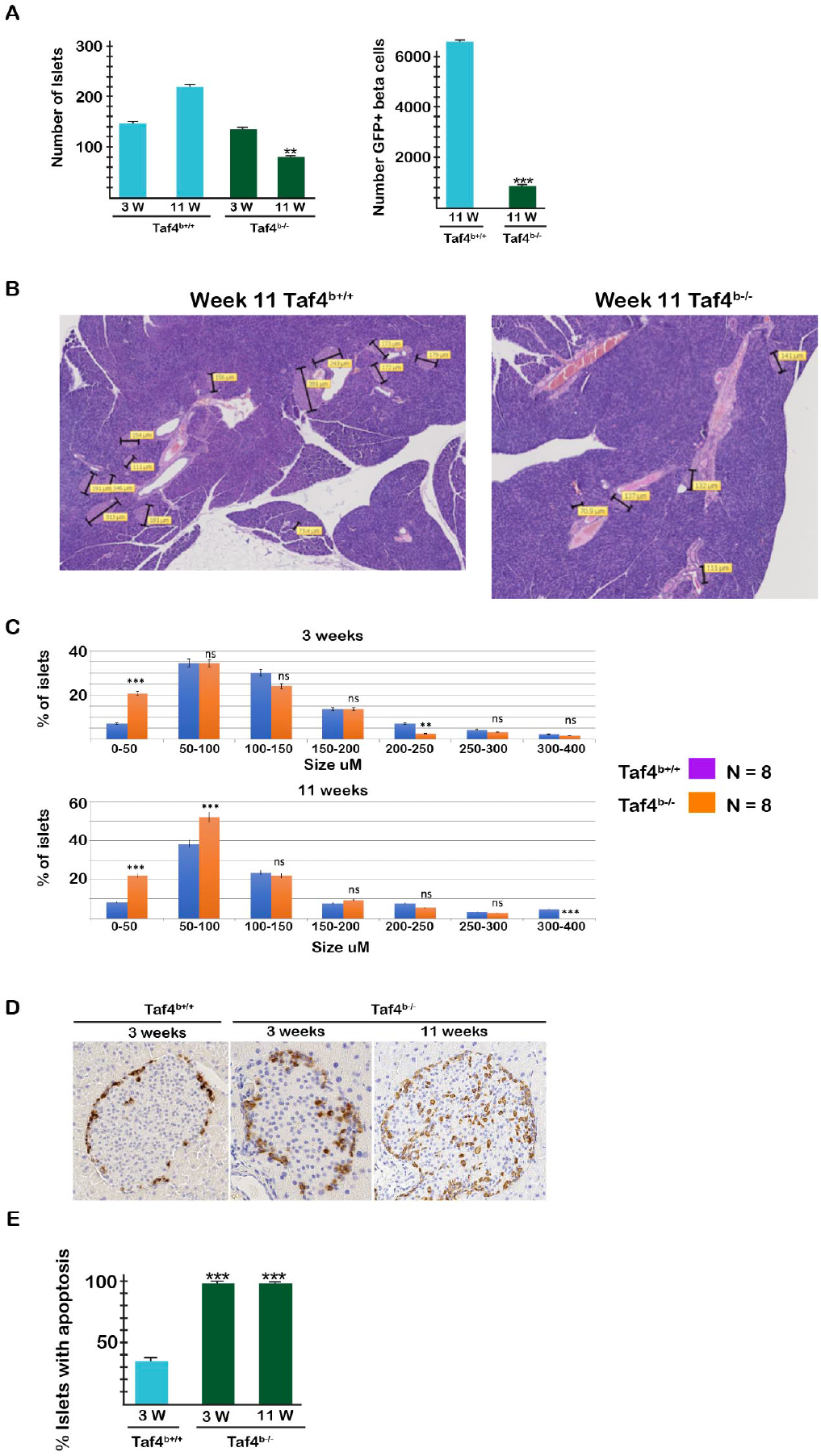
Long term fate of BCs following Taf4 inactivation. **A.** Number of islets in *Taf4*^*b−/−*^ and control animals. **B.** Histology of pancreas sections used to measure islet size distribution in *Taf4*^*b−/−*^ and control animals in panel **C**. **D.** Tunel staining of islets from in *Taf4*^*b−/−*^ and control animals. **E**. % counted of islets with apoptotic BCs. Student T-test ** p<0.05, *** p<0.001 from N=8.

Examination of BC histology by electron microscopy showed numerous densely stained mature insulin granules in the cytoplasm in control BCs (Fig. S1A). However, after 2-3 weeks, the number of mature insulin granules was strongly diminished and replaced by lighter staining pro-insulin granules (Fig. S1B-C). Furthermore, the Golgi apparatus was visibly enlarged in mutant BCs. After 11 weeks two BC populations could be discerned. Around 70% of BCs displayed only few mature insulin granules in their cytoplasm (Fig. S1D), whereas the other 30% displayed more numerous granules (Fig. S1E). Increased extracellular space between the mutant BCs was observed compared to the tight cell-cell contacts of control BCs (Fig. S1F).

Together the above results showed that loss of Taf4 in BCs led to both acute and longer-term effects. Within 1-3 weeks, diminished insulin maturation was observed along with defective GSIS. As a consequence, mutant animals showed persistent hyperglycemia. Over the next weeks, the number of BCs was reduced due to apoptosis, but not all BCs were lost and surviving BCs with mature insulin granules or predominantly pro-insulin granules were observed at later times accounting for stabilised blood glucose and insulin levels and long-term survival. Moreover, by 34-53 weeks, islets were partially repopulated by Taf4-expressing BCs. Thus, after an initial acute effect of Taf4 loss on the existing BC population, the surviving BCs adapted and partially restored their insulin signalling capacity and at later times were partially replaced by a new Taf4-expressing BC population.

### Taf4 regulated gene expression programs in beta cells

To investigate how Taf4 knockout in BCs impacted gene expression as the basis for the observed phenotypes, we performed RNA-seq from WT islets and from islets, 1, 3 and 5 weeks after Tam injection (Fig. 4A-B and Dataset S1). Already 1 week after Tam injection, more than 600 genes were down-regulated and 390 up-regulated (Figs. 4A and S2A-B). At 3 and 5 weeks, the number of de-regulated genes strongly increased such that by 5 weeks more than 2500 genes were down-regulated and 900 up-regulated (Fig. S2A-B). More than 300 genes were down-regulated under all conditions together with a large overlap of more than 800 between weeks 3 and 5. At 1 week, several genes critical for BC function such as *Slc2a2*, *Trpm5*, *Ins1* and genes involved in calcium signalling and cell contact were down regulated (Fig. S2C). In contrast, up-regulated genes were strongly enriched in inflammatory/stress response including numerous chemokines and *Reg3a*, *Reg3b* and *Reg3g*, genes with important roles in islet stress response and regeneration (Fig. S2C). Up regulated genes were also strongly enriched for oxidative phosphorylation and DNA metabolism including repair, replication and general transcription factors (Fig. 4A and Dataset S1). Expression of the AC and DC markers *Gcg* and *Sst* was also rapidly up-regulated and then attenuated with time (Fig. S2C).

**Figure 4.**
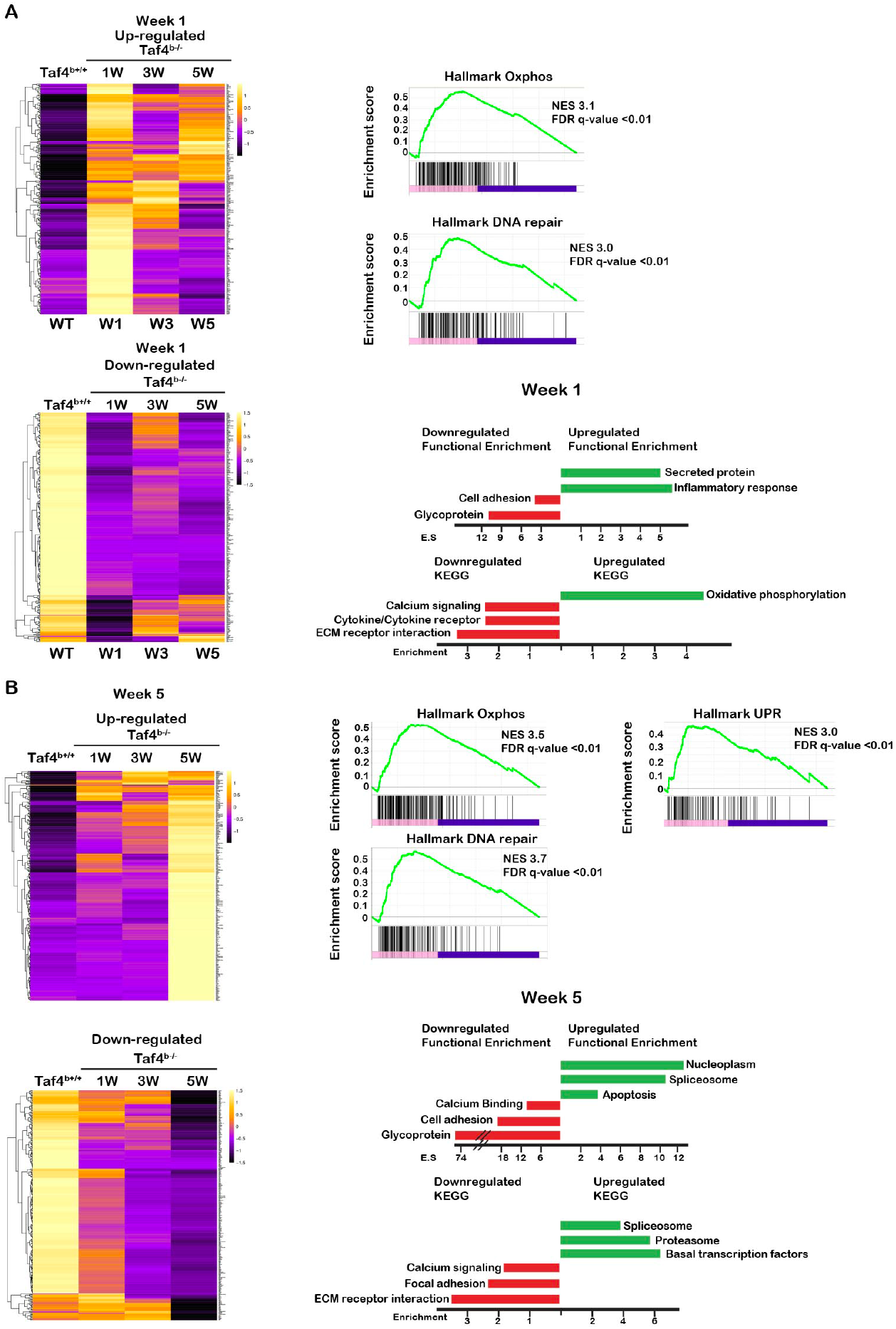
Effects of Taf4 inactivation on gene expression of isolated islets. **A.** Heatmap showing the expression of the 100 genes most up and down-regulated 1 week after Taf4 inactivation. Right hand panels show the GSEA and ontology analyses of the de-regulated genes. **B.** Heatmap showing the expression of the 100 genes most up and down-regulated 5 weeks after Taf4 inactivation. Right hand panels show the GSEA and ontology analyses of the de-regulated genes.

While overall an increasing number of genes was deregulated with time, we identified sets of genes whose normal expression was partially restored. Expression of a subset of genes strongly down-regulated after 1 week was progressively restored to more normal levels after 3 and 5 weeks (Figs. 4A and S2D). Similar results were seen with up-regulated genes the most striking example of which are the inflammatory/stress response genes up-regulated after 1 week but restored to almost normal after 3 and 5 weeks.

After 5 weeks, down regulated genes remained enriched in those involved in cell adhesion and calcium signalling, many of which were already de-regulated at 1 and/or 3 weeks (Fig. 4B). Similarly, genes involved in oxidative phosphorylation and DNA metabolism remained up-regulated along with novel pathways such as the unfolded protein response (UPR). In addition, genes involved in diverse aspects of RNA metabolism were also up-regulated along with a collection of basal transcription factors and apoptotic genes in keeping with the increased apoptosis described above. Down-regulated genes additionally comprised components of multiple signalling pathways such as FGF, SMAD/TGFB and Notch (Dataset S1).

Up-regulation of genes involved in chromatin organisation, RNA metabolic functions and transcription were already observed after 3 weeks (Fig. S2D). These observations suggested that the initial stress response immediately after Taf4 inactivation was followed by chromatin reorganisation and compensation for loss of *Taf4* by up-regulation of *Taf4b* and *Taf12*. Indeed, *Taf4b* expression was much lower than that of *Taf4* in wild-type islets. Note that as previously described, the deletion of the C-terminal Taf4 exons had little effect on *Taf4* mRNA levels and thus *Taf4* did not appear as a downregulated transcript.

We assessed the effect of Taf4 inactivation on chromatin accessibility using ATAC-seq on isolated islets. After 1 week, extensive changes in chromatin accessibility were observed with a global increase in accessibility upon Taf4 loss. When focussing on the proximal promoter, the number of sites enriched in the WT was higher than those enriched in the mutant (Fig. S3A). A similar result was seen after 5 weeks, where again more promoter proximal sites were enriched in the WT than in the mutant (Fig. S3B). To identify transcription factors whose occupancy was modified, we carried out *in silico* footprinting analyses of the differentially accessible regions at the proximal promoter ^39^. Binding of Hox-domain containing transcription factors including critical BC factors such as Nkx6-1 and Hnf1b were enriched in the WT, whereas factors like Yy2, Arnt2, Bhlhe41 and Mycn were enriched 1 week after Taf4 loss (Fig. S3C). Similarly, after 5 weeks, enriched binding of several factors including Yy2, Nrf1 was observed (Fig. S3D).

Together the ATAC-seq and RNA-seq data indicated that Taf4 loss led to a reduction in chromatin accessibility at proximal promoters, modified transcription factor binding with reduced binding of critical BC factors together with reduced expression of a collection of genes essential for BC function and enhanced binding of a new set of transcription factors.

### Taf4 inactivation results in loss of beta cell identity and trans-differentiation into alpha-like cells

While the above data provided an overall view of gene expression changes in islets, they did not assess whether Taf4 loss affected only BC gene expression or if it impacted gene expression in other cell types as suggested by increased *Gcg* and *Sst* expression. To address this, we performed single cell (sc)RNA-seq from purified dissociated Langerhans islets from control Taf4-expressing mice and *Taf4*^*b−/−*^ mice 1 and 5 weeks after Tam injection.

Analyses of around 12000 cells from Taf4-expressing control islets identified the major endocrine populations, with 4 BC clusters expressing high levels of *Ins1* and *Ins2* (clusters 0-2 and 4 in Fig. 5A). Cluster 3 represented *Gcg*-expressing ACs, cluster 6 *Sst*-expressing DCs and cluster 5 an acinar population (Fig. 5A). The 4 BC sub-populations could be distinguished by their expression signatures with a strong ER stress signature in cluster 2, while cluster 0 represented potential secretory cells with high expression of *Slc2a2*, *Abcc8* and *G6pc2* and lower *Ins1*/*2* expression that was more prominent in clusters 1 and 4 (Fig. S4).

**Figure 5.**
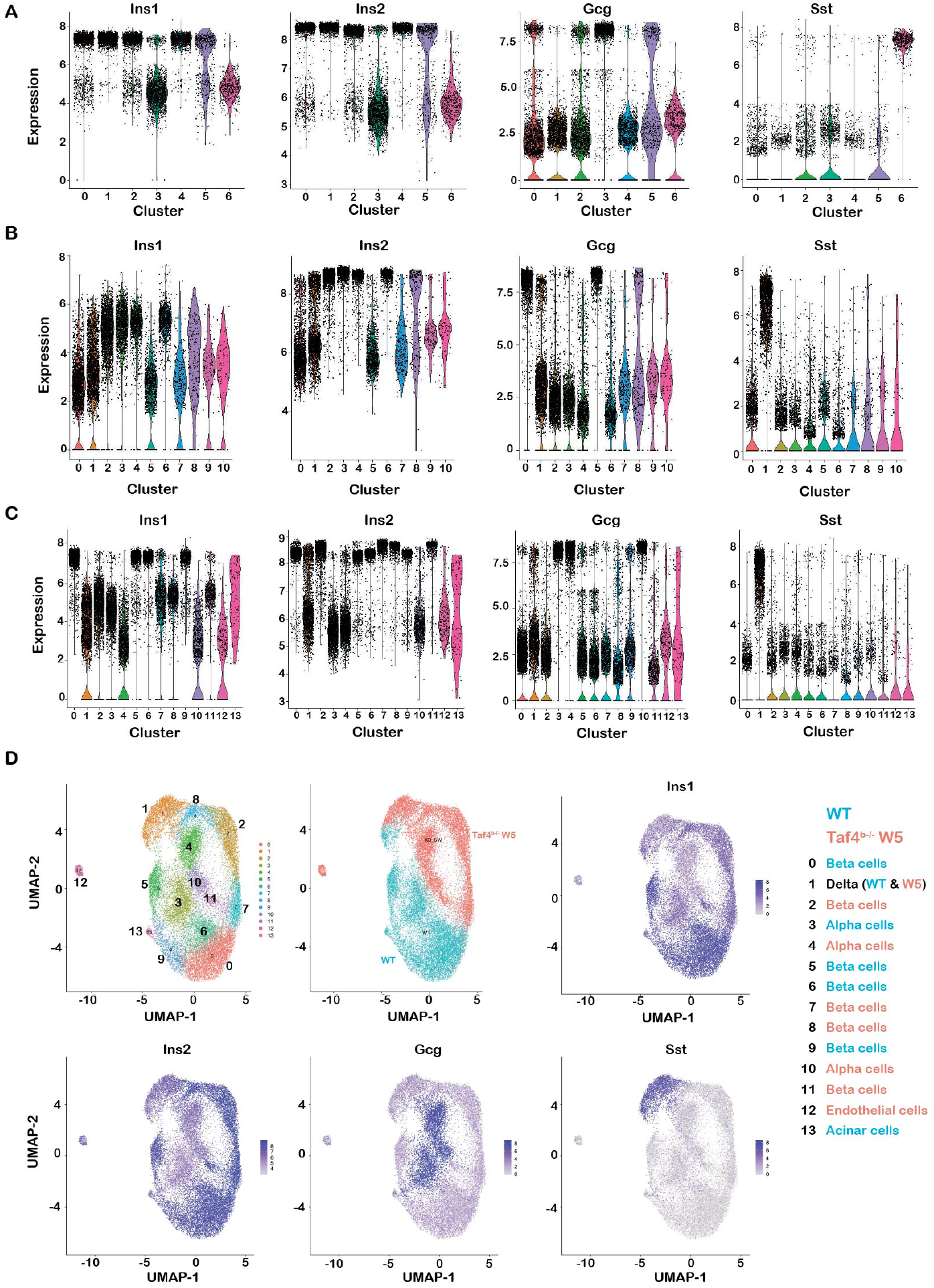
Sc-RNA-seq of WT and week 5 mutant islets. **A-C.** Violin plots of expression of the indicated genes in each cluster from WT islets (A), 5 week mutant islets (B) and the aggregate (C). **D**. UMAP representations of the aggregate data, illustrating origin of cells from WT and mutant animals and the expression of the genes indicated in each panel.

We analyzed 10 666 cells 5 weeks after Taf4 inactivation distinguishing again 4 BC populations with high *Ins2* expression (clusters 2-4 and 6, Fig. 5B), but reduced *Ins1* expression as noted in the bulk RNA-seq. Cluster 1 corresponded to *Sst*-expressing DCs, while clusters 0 and 5 corresponded to *Gcg*-expressing ACs. We aggregated the WT and week 5 data sets allowing a global comparison of all cell populations under the 2 conditions (Figs. 5C, D and S5) clearly distinguishing BC populations from WT and mutant conditions based on high *Ins1* expression in WT (clusters 0, 5, 6 and 9) from the *Ins2* high/*Ins1* low mutant populations (clusters 2, 7, 8 and 11). The WT and mutant cell populations segregated in a UMAP representation (Fig. 5D) where WT and mutant BCs were distinguished by strongly reduced expression of the BC markers *Ucn3* and *Slc2a2* as seen in bulk RNA-seq (Fig. S6A). High expression of stress markers *Ddit3* and *Herpud1* characterized WT BCs in cluster 5 and mutant clusters 2 and 8 (Fig. S6A). The aggregate also revealed 3 distinct AC populations, cluster 3 from WT and clusters 4 and 10 from the mutant with a strong stress response signature in cluster 4 (Fig. S5). DCs from the two conditions adopted a more homogenous population (cluster 3).

The important changes in gene expression following Taf4 inactivation in BCs allowed clear distinction between the WT and mutant BC populations and segregation of AC populations from WT and mutant with 2 populations in the mutant. DC gene expression on the other hand was less affected with little difference in the populations of the two conditions.

We analyzed 2295 cells 1 week after Taf4 inactivation. Clusters 2 and 5 corresponded to the most differentiated BC populations with high expression of *Ins2* and lowest levels of *Gcg* and *Sst* as well as high levels of the mature BC marker *Ucn3* (Fig. 6A). Cluster 2 however was marked by ER stress factors (Fig. S6D). Cluster 0 showed higher *Gcg* and *Sst* expression, although lower than in differentiated AC and DC populations, clusters 3 and 7, respectively, that segregated from the BC populations in the UMAP and tSNE representations (Figs. 6B-C and S6B). The small cluster 6 also showed characteristics of a mixed identity population expressing *Ins1*/*2* and intermediate levels of *Gcg*. Taf4 inactivation therefore led to appearance of a large population with mixed identities expressing *Ins1* and *Ins2*, but also *Gcg* or *Sst* and sometimes all three.

**Figure 6.**
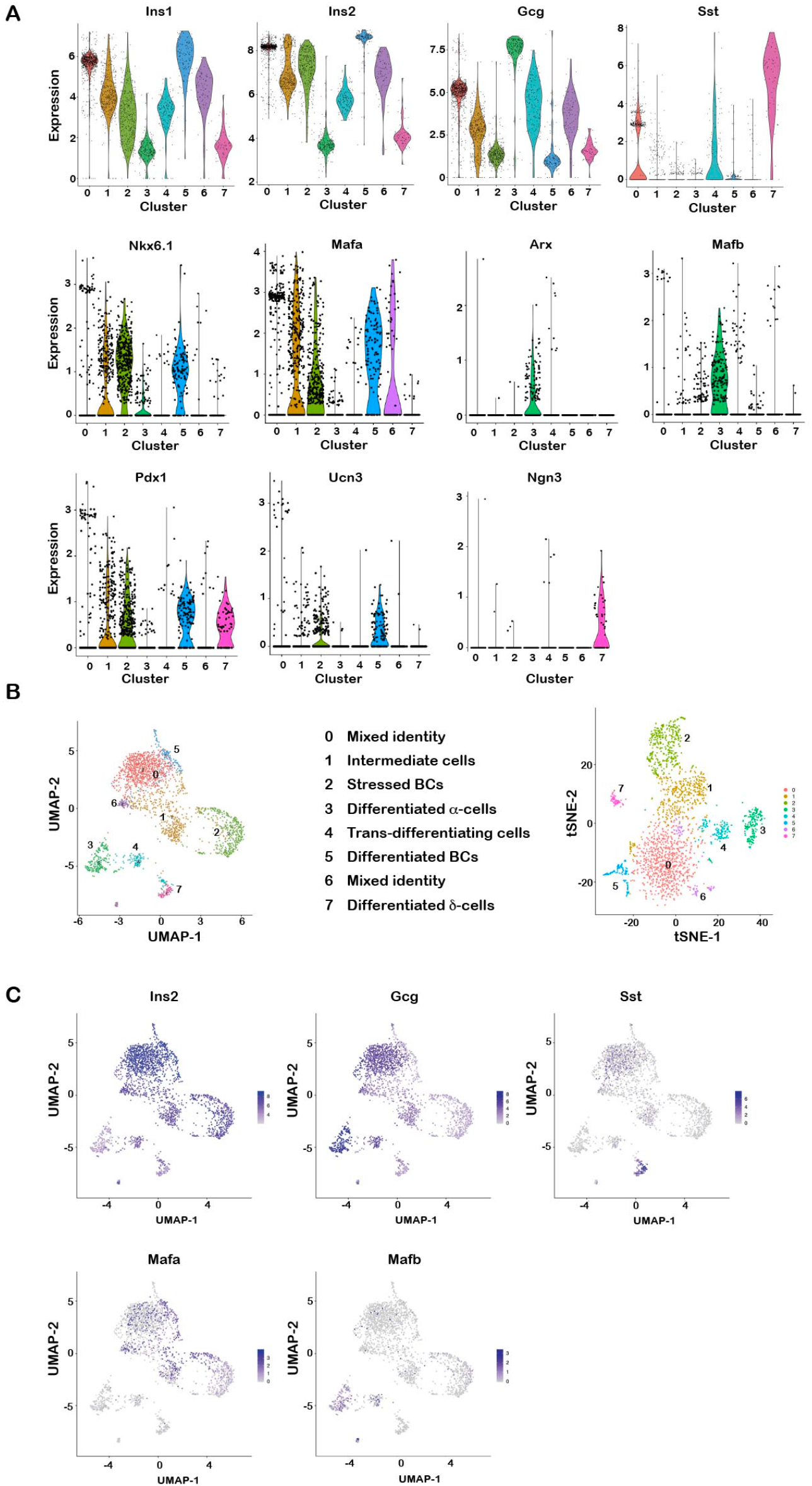
Sc-RNA-seq of week 1 mutant islets. **A.** Violin plots of expression of the indicated genes in each cluster. **B**. UMAP (left) and tSNE (right) representations of the cell populations. **C.** UMAPs illustrating the expression of the genes indicated in each panel.

Cluster 1 appeared to be intermediate cells with lower *Ins2* and higher *Gcg* expression than the more differentiated clusters 2 and 5 and lowered expression of the mature BC marker *Ucn3*. Cluster 4 expressed intermediate levels of *Ins2*, *Gcg* and *Sst* and segregated between the BC, AC and DC populations on the UMAP and between the BC and AC populations in the tSNE. Cluster 4 cells lost expression of critical BC-markers such as *Pdx1*, *Mafa* and *Nkx6.1* and appeared to be trans-differentiating intermediates between BC and AC-like and perhaps also DC-like identities (Figs. 6C and S6B). This was further supported by the ontology of the differentially regulated genes (Figs. S6C and Dataset S2). Cluster 5 was marked by differentiated BC markers, cluster 2 showed translational activity, ER stress associated with high *Ddit3* and *Herpud1* expression and high oxidative phosphorylation signatures (Fig. S6D). In contrast, signatures for transcription regulation, chromatin modification and RNA metabolism were hallmarks of clusters 1 and 4 consistent with the idea that these cells were undergoing transcriptional reprogramming.

We used the SCENIC program ^42^ to identify transcription regulatory networks active in the different cell populations. The SCENIC-based tSNE (Fig. 7A) that groups cells based on their regulon activities differed from the expression based tSNE (Fig. 6B). With SCENIC, the BCs clusters 2 and 5 grouped together, clusters 1 and 4 grouped closely, while cluster 0 grouped close to ACs (Fig. 7A). Consistent with the idea that cluster 5 were differentiated BCs, they showed high activity of the Mafa, Nkx6-1 and Neurod1 regulons (Fig. 7B-C). These regulons were active in cell sub-populations of clusters 0 and 1, but absent from ACs and DCs. Cluster 2 was marked by activity of Ddit3 and Atf5, that promotes BC cell survival under stress ^43^, consistent with the elevated stress response in this population (Fig. 7C-D). In contrast, Mafb regulon activity was seen in ACs and in clusters 0, 1 and 4 confirming they represented abnormal populations with mixed identities, with some cells showing both Mafa and Mafb activities (Fig. 7B-C). Transition from cluster 1 to cluster 4 was therefore marked by a switch away from BC identity factors to the AC determinant Mafb.

**Figure 7.**
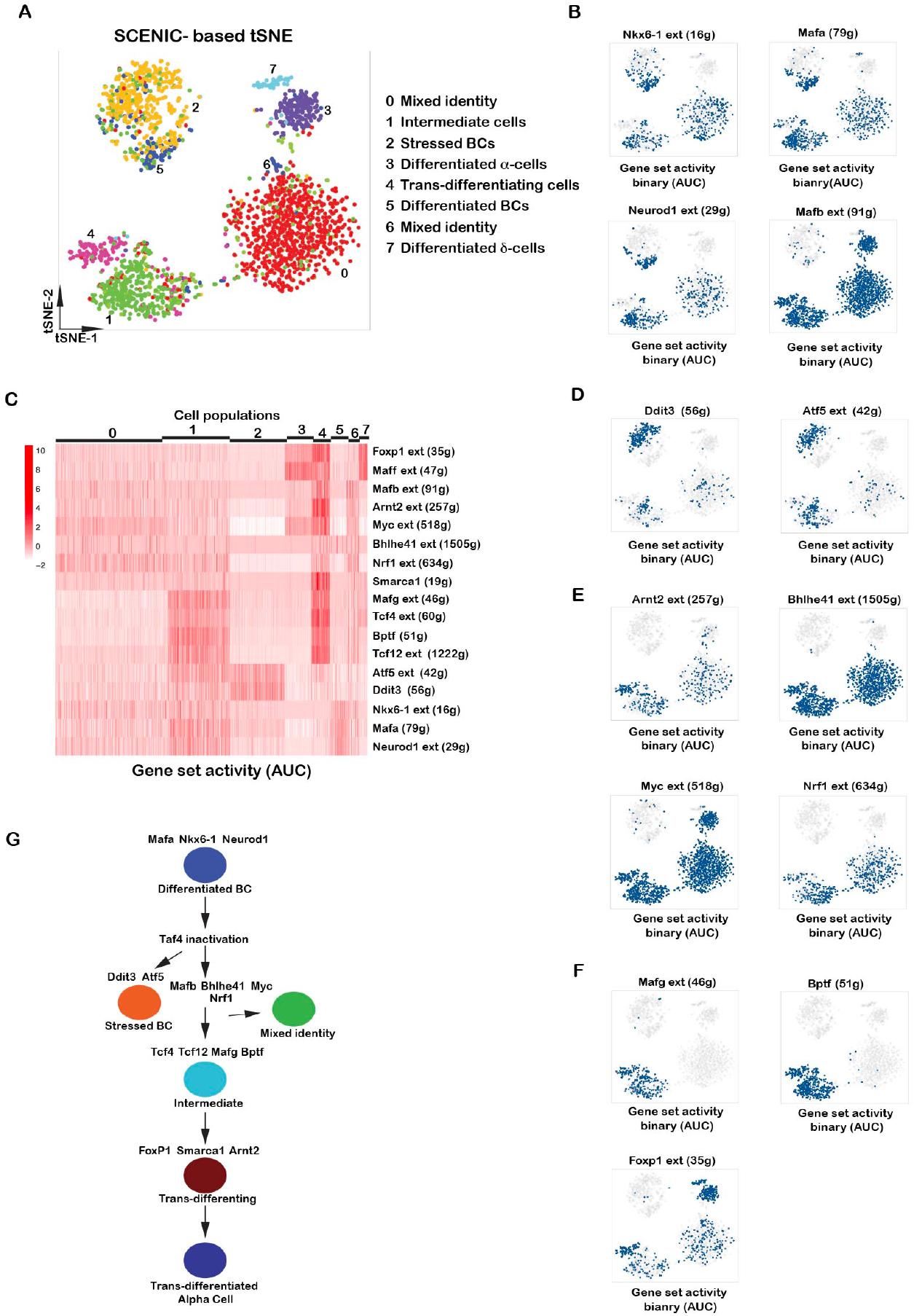
Transcription factors regulating cell state in week 1 mutant islets. **A**. SCENIC-based tSNE representation coloring cells based on the binary activities of the transcription factor regulons. **B, D-E**. Binary activities of the transcription factor regulons in the different cell populations. **C.** Heatmap of regulon activities in the different cell populations were quantified using AUCell. **G.** A flow-chart of the progression of cell state from differentiated BCs to trans-differentiated alpha cells. The SCENIC regulons marking the transitions of each state are indicated.

Strikingly, clusters 0, 1 and 4 showed activity of the Arnt2, Bhlhe41, Myc and Nrf1 regulons corresponding to enhanced binding predicted from ATAC-seq footprinting (Fig. 7C and E). The abnormal cell populations were therefore marked by activity of regulons not active in more differentiated BCs (clusters 2 and 5). Clusters 1 and 4 were specifically marked by high activity of the small Maf inhibitor Mafg, Tcf4 and Tcf12 as well as the Bptf subunit of the NuRF complex coherent with the chromatin remodeling ontology signature of this cluster (Fig. 7C and E). The Foxp1 and Maff regulons showed highest activity in cluster 4 and in differentiated alpha and/or delta cells (Fig. 7C and F). Mafb, Myc, Arnt2, Mafg, Tcf4, Tcf12, Bptf and Foxp1 regulons showed highest combinatorial activity in cluster 4. Moreover, activity of Smarca1, the catalytic subunit of the NuRF complex, was specific for cluster 4 that together with Bptf activity pointed to a critical role of the NuRF complex in this cluster (Fig. 7C). These results were consistent with a major transcriptional reprogramming and chromatin remodeling associated with trans-differentiation. As a result, by 5 weeks two AC populations were observed potentially corresponding to the normal and trans-differentiated populations. We also noted that trans-differentiation did not appear to involve transition via a precursor state as low *Ngn3* (Neurog3) expression was seen in the DC population, but was not significantly expressed in clusters 1 and 4 (Fig. 6A).

At 5 weeks, no populations with mixed or obviously trans-differentiating characteristics were observed suggesting that trans-differentiation took place immediately after Taf4 loss and was terminated by 5 weeks. To assess this in more detail, we performed SCENIC analyses on 2500 randomly selected cells from the WT/W5 aggregate. The SCENIC based tSNE again grouped cells in a manner different from the expression based tSNE (Figs. 5D and S7A). Regulome analyses showed that WT BCs were principally marked by Mafa and the glucose sensing transcription factor Mlxipl that was high in the WT BC populations 0 and 6 (Fig. S7B). Mafa activity was also strong in the mutant BC populations with cluster 11 displaying high activity of both Mafa and Mlxipl. Mafb activity was high in the three AC populations, but contrary to week 1 no overlap of Mafa and Mafb activities was seen. Several mutant BC populations characterized with stress response as described above did however show activity of the Mafg, Yy1, Atf5, Atf3, and Bhlhe41 regulons together with those of Jun and Fosb (AP1). Similarly, the stressed mutant ACs also showed elevated AP1 and Atf3 activity. Mutant BCs also showed elevated activity of chromatin remodeling factor Chd2. In contrast, activity of all of these regulons was much lower or absent in WT BCs. These data showed that WT and mutant BCs and ACs showed different regulon activities with elevated activity of several factors involved in stress response in the mutant populations.

To verify the idea that a sub-population of BCs trans-differentiate into Gcg-expressing apha-like cells, we performed immunostaining with antibodies against Ins and Gcg (Fig. S8). In WT islets, Gcg staining was limited to ACs surrounding the BCs within the islet. Strikingly, by 3 weeks after Taf4 inactivation, Gcg-expressing cells were observed both around and within the islets amongst the Ins-expressing BCs, a phenomenon that was seen at all subsequent stages until 55 weeks after injection and reminiscent of what was seen upon loss of Pdx1 that promoted to BC trans-differentiation into Gcg-expressing cells ^44^. Such cells were not co-stained with Ins, but were labelled uniquely by Gcg. Together with the results of the sc-RNA-seq, these observations support the idea that a sub-population of BCs trans-differentiate into a novel population of Gcg-expressing cells within the islets.

## Discussion

### Taf4 is essential for normal beta cell function

We show that Taf4 inactivation in adult murine BCs led to increased glycaemia due to defective insulin signalling and secretion, a consequence of an immediate and major impact on gene expression in BCs and at later times apoptosis of part of the BC population. Despite this, the surviving BCs allowed stabilization of glycaemia and longer-term survival of the animals.

Immunostaining showed a rapid and efficient loss of Taf4 expression in BCs by 1 week after Tam injection. At 34 weeks, Taf4-expressing BCs were again detected, but their proportion did not increase by 55 weeks. While it is possible that Taf4-expressing BCs detected at later times arose from rare non-recombined BCs, no Taf4 expressing BCs were observed upon staining of multiple islets at regular intervals from 3-24 weeks. Furthermore, the number of Taf4-expressing BCs did not increase between 34 and 55 weeks arguing against proliferation of non-recombined BCs. Alternatively, Taf4-expressing BCs may arise from trans-differentiation of ACs or other islet populations into BCs as has been previously described, for example under conditions of reduced BC-mass ^2, 5, 6, 12, 45^, one of the consequences of Taf4 inactivation. Notwithstanding the mechanisms involved, these data showed that despite reduced BC mass through increased apoptosis and an ensuing elevated glycaemia, the BC population was only partially and slowly replaced by newly generated BCs.

Taf4 inactivation led to a long-term reduction in BC mass and reduced islet size and by 5 weeks, a majority of BCs were filled with pro-insulin granules rather than mature insulin granules. Despite these observations and long-term reduced expression of key BC function genes (*Slc2a2*, *Ins1*, *G6pc2, Trpm5*), glycaemia was stabilized and even mildly reduced by 10 weeks, well before the first Taf4-expressing BCs were observed at 34 weeks. Indeed, bulk RNA-seq identified sets of genes whose expression was partially corrected at 5 weeks along with up-regulation of *Taf4b* and its heterodimerization partner *Taf12,* a potential compensatory response to Taf4 loss. Moreover, only a sub-population BCs underwent apoptosis, while others displayed long-term survival. This is perhaps related to BC heterogeneity, as WT BCs with high and low levels of ER stress markers were identified consistent with previous reports ^9, 46, 47, 48, 49, 50^. ER stress is modulated by insulin demand and has been associated with BC differentiation, survival and proliferation. Perhaps differences in ER-stress amongst BC populations contributed to their ^10, 50^. Furthermore, while *Ins1* expression was diminished, *Ins2* remained strongly expressed in mutant BCs and although insulin maturation was diminished it was not completely lost. In addition, proinsulin is secreted and retains a some of the metabolic activity of mature insulin. Thus, stabilization of an albeit elevated glycaemia compatible with long-term survival of the Taf4^b−/−^ animals can be can be accounted for by persistence of partially functional BCs for many weeks before the appearance of new Taf4 expressing BCs.

### A model for beta cell trans-differentiation

Bulk RNA-seq and ATAC-seq showed that Taf4 inactivation had a potent impact on chromatin accessibility and gene expression with diminished binding of factors important for BC identify. In contrast, increased binding of several transcription factors was predicted from ATAC-seq and several of the corresponding regulons were activated in the abnormal cell populations after 1 week and in subpopulations of mutant BC and AC at 5 weeks. At week 1, cells with mixed identities displayed regulon activity of BC factors, of the AC determinant Mafb and a combination of several others factors. These cells did not express *Arx* and therefore displayed an incomplete switch in identity in line with the intermediate levels of *Gcg* compared to fully differentiated ACs. Previous studies on normal mouse or human pancreas by immune-staining or sc-RNA-seq detected only rare cells simultaneously expressing two of the islet hormones ^9, 10, 46, 47, 48, 49^. Such cells have however been detected in pathological situations like type 1 and type 2 diabetes and in mice following loss of Arx and Dnmt1 ^9^. The appearance of a population of mixed identity cells was therefore a consequence of Taf4 inactivation and potentially the resulting activation of novel regulons.

Our data pointed to a process of stress-induced BC trans-differentiation. Cluster 1 cells maintained activity of BC factors, but acquired activity of Mafb and a combination of other factors. Cluster 4 cells displayed strongly decreased expression and activity of BC factors, but showed Mafb activity, increased *Arx* and higher *Gcg* and *Sst* expression. Furthermore, cluster 4 cells were strongly marked by Foxp1 activity consistent with the reported essential role of Foxp factors in post-natal ACs ^51^ and by activity of Maff, Mafg and the NuRF complex. Inhibition of (s)Maf proteins in BCs led to increased Mafa activity and BC function correlating with the increased Mafg/Maff activity and loss of BC identity in the trans-differentiating cells found here ^52, 53^. Nevertheless, the molecular details of sMaf function in regulating Mafa, Mafb in pancreas remain to be fully addressed.

We propose a model for BC trans-differentiation involving a progressive loss of BC identity associated with gain of activity for Myc, Mafg, Arnt2, Bhlhe41, Tcf4, Tcf12, Nrf1 and Mafb (Fig. 7G). A population of cells further undergoes chromatin remodelling and transcriptional reprogramming through the combinatorial action of these factors and subsequently acquisition of activity of Foxp1, Maff and Smarca1 that promotes trans-differentiation into *Gcg*-expressing alpha-like cells and perhaps also delta-like cells. The appearance of trans-differentiated alpha-like cells was further attested by two distinguishable AC populations after 5 weeks and by the appearance of Gcg-expressing cells intermixed amongst BCs, a phenomenon not seen in WT islets. The fact that no Taf4 expression was seen in islets at this time, further supports the idea that these Gcg-expressing cells are derived from Taf4-null BCs.

The mechanisms of trans-differentiation of ACs to beta-like cells have been documented ^4, 8, 9: Thorel, 2010 #9248, 54^, and examples of BC to alpha-like cell trans-differentiation often involve genetically induced loss of BC determinants ^11, 14, 44^. In this study, BC trans-differentiation also involved loss of expression and activity of critical BC-determinants that may be triggered by the potent, but transient stress and inflammatory response with strong expression of several Reg-family proteins and multiple chemokines seen 1 week after Taf4 inactivation. This inflammatory/stress environment may be responsible for activation of Myc activity and previously implicated in BC de-differentiation ^50, 55, 56^.

Previous studies of trans-differentiation often focussed on transcription regulators with known roles in BC and AC identity. However, BC cell populations actively undergoing trans-differentiation and the associated transcriptional regulatory programs involved have been only poorly characterized. Using an un-supervised bioinformatics approach, we defined how novel combinations of known critical BC and AC determinants act in combination with additional transcription and chromatin remodelling factors to promote BC trans-differentiation. While different cues may underlie initiation of BC trans-differentiation in pathological situations or following Taf4 inactivation, further studies will determine whether they share common aspects of the transcriptional reprogramming described here.

## Acknowledgements

We thank, I Michel for excellent technical assistance, all the staff of the IGBMC common facilities in particular the IGBMC animal, high throughput sequencing (GenomEast) and electron microscopy facilities, the Mouse Clinical Institute for help with measurement of insulin and glucose levels. This work was supported by institutional grants from the Centre National de la Recherche Scientifique, the Institut National de Santé et de la Recherche Médicale, the Université de Strasbourg, the Ligue Nationale contre le Cancer, Agence Nationale de la Recherche (ANR)-15-CE14-003 project Dysmetaf, the ANR-10-LABX-0030-INRT French state fund under the framwork programme Investissements d’Avenir ANR-10-IDEX-0002-02. The IGBMC high throughput sequencing facility is a member of the “France Génomique” consortium (ANR10-INBS-09-08). ID is an ‘équipe labellisée’ of the Ligue Nationale contre le Cancer. TK was supported by fellowships from the ANR and the FRM. All sequencing data reported here have been submitted to the GEO database under accession number GSE151366.

## Author contributions

TK performed the physiological, histological and gene expression analyses, GD performed bioinformatics analyses, IM performed immunostaining, GM managed the breeding, housing and genotyping of the mice. TK, GD and ID conceived the experiments analyzed the data and wrote the paper.

---

**Supplemental Figure 1.**
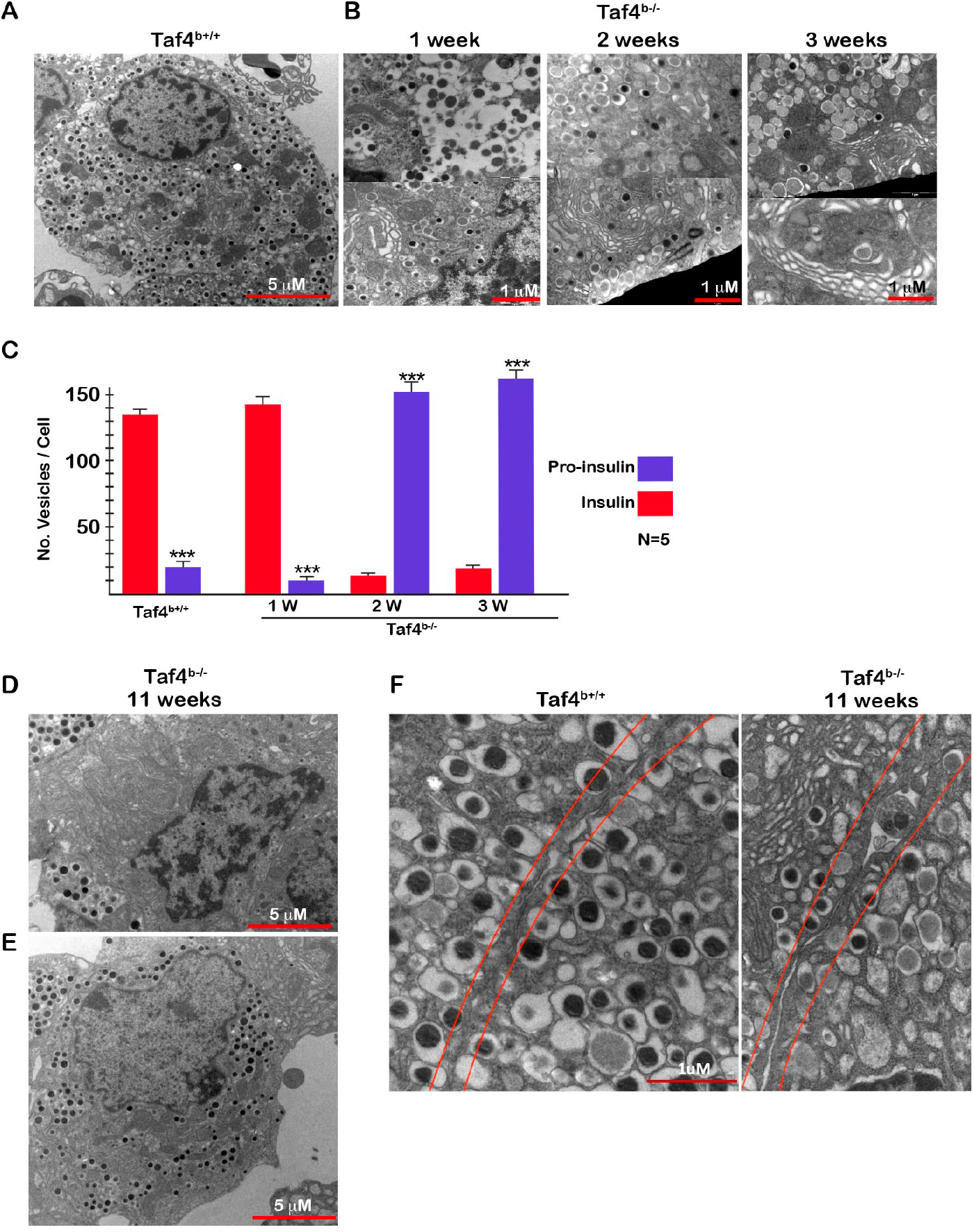
Ultrastructure of BCs by electron microscopy. **A**. BC from control animal with densely staining insulin secretory granules in the cytoplasm. **B**. BCs from *Taf4*^*b−/−*^ animals highlighting accumulation of lighter staining proinsulin secretory granules and expanded Golgi apparatus. **C.** Quantitation of insulin and proinsulin secretory granules. **D-E.** BCs from *Taf4*^*b−/−*^ animals 11 weeks after Tam injection illustrating cells with few (70%, panel D) or more numerous (30% panel E) insulin secretory granules. **F.** Control and mutant BCs highlighting the cell-cell contacts and the loss of tight contacts between the mutant cells. Student T-test *** p<0.001

**Supplemental Figure 2.**
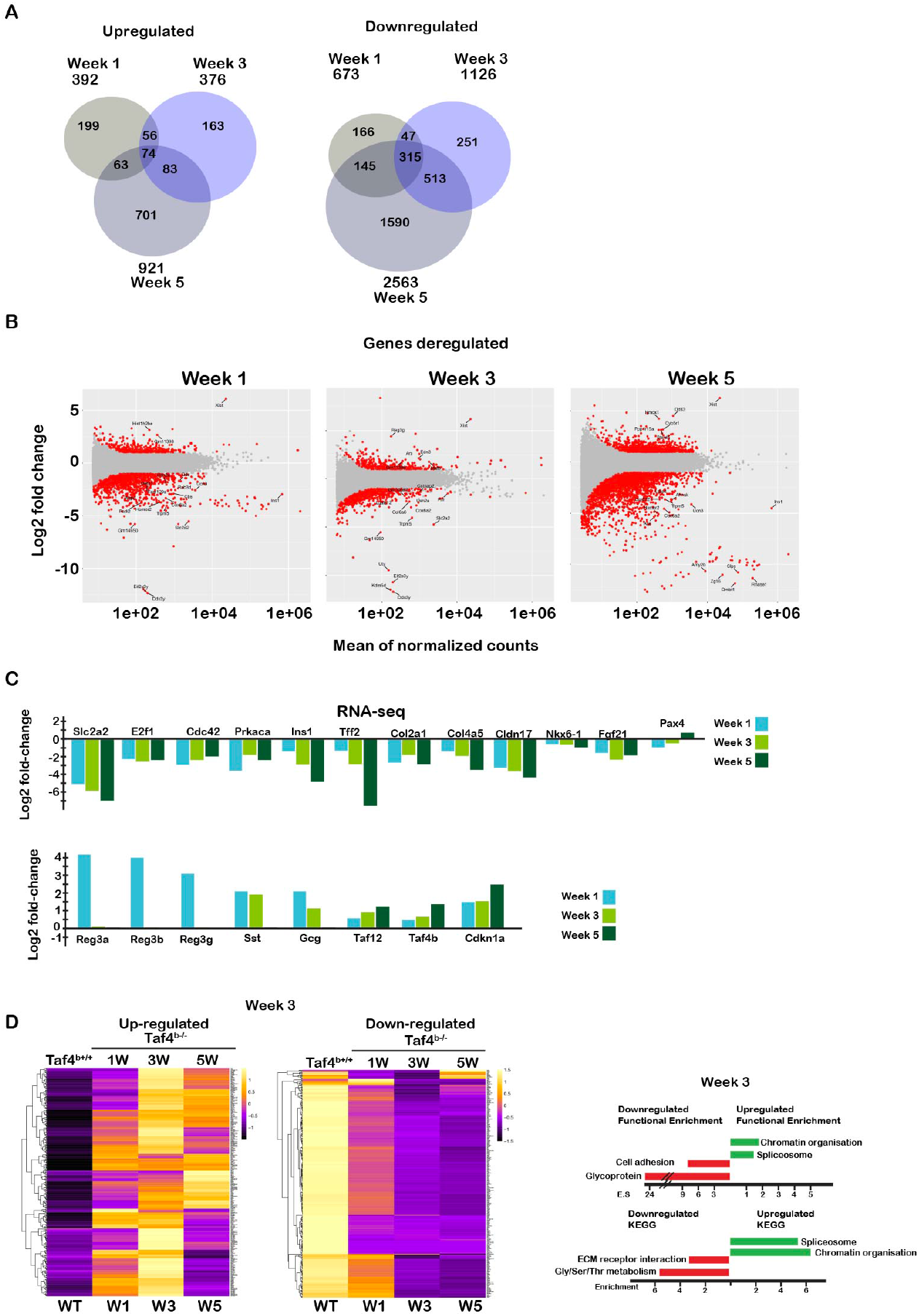
Changes in gene expression upon Taf4 inactivation. **A.** Venn diagrams summarizing genes up and down-regulated 1, 3 and 5 weeks after Tam injection. **B**. Volcano plots showing changes in gene expression after 1, 3 and 5 weeks. **C.** Summary of RNA-seq data on the expression of selected up and down-regulated genes critical for BC function, identity, cell-cell contact, stress response and AC and DC markers. **D**. Heatmap showing the expression of the 100 genes most up and down-regulated 3 weeks after Taf4 inactivation. Right hand panel shows ontology analyses of the de-regulated genes.

**Supplemental Figure 3.**
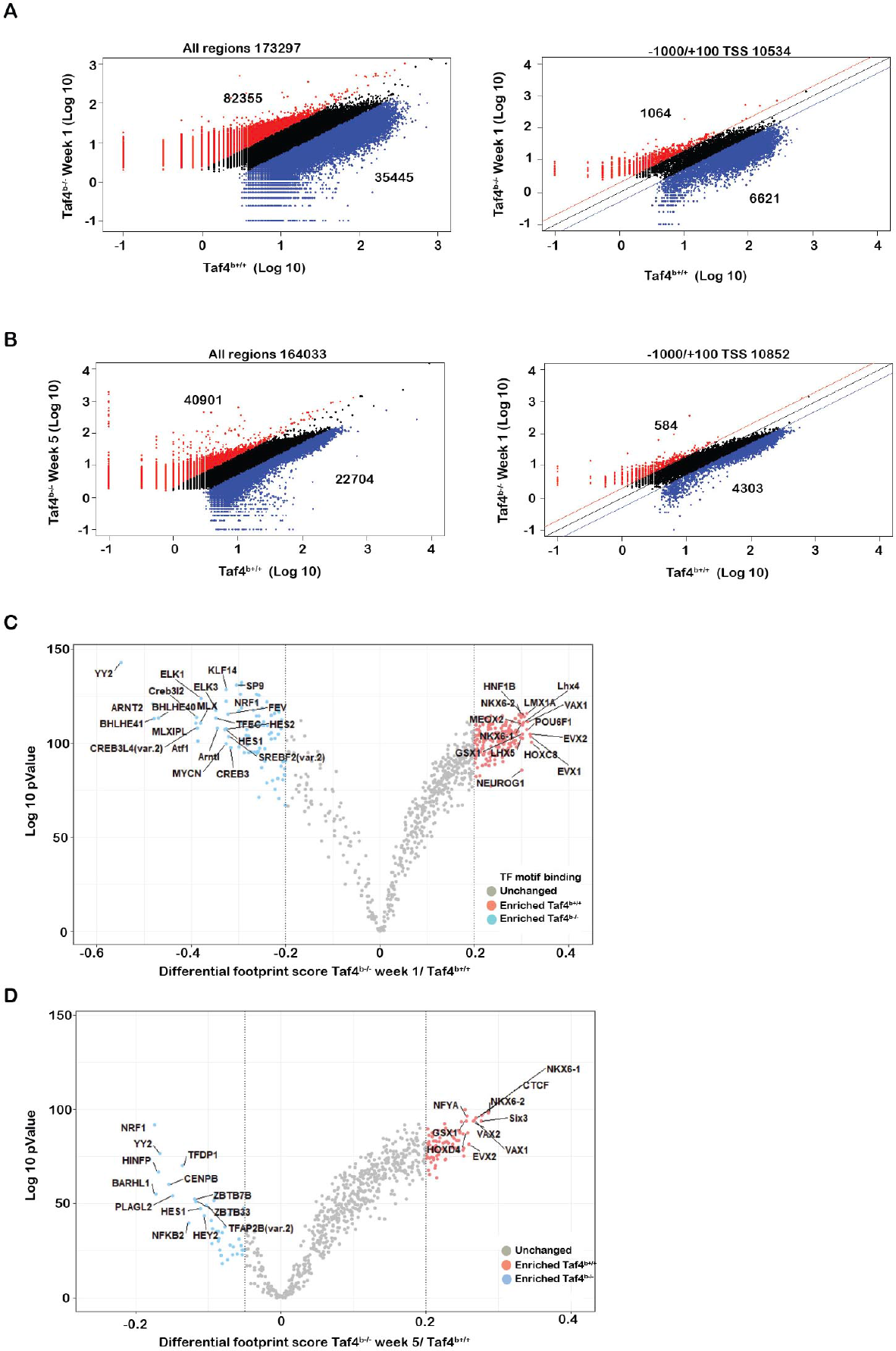
Changes in chromatin accessibility in isolated islets 1 and 5 weeks after Taf4 inactivation measures by ATAC-seq. **A-B**. Scatterplots showing Log10 changes in ATAC-seq peaks either globally (left panels) or at the proximal promoter (−1000/+100 base pairs relative the transcription start site, TSS, right panel) at 1 week (A) or 5 weeks (B) following TAF4 inactivation. The number of peaks enriched in each condition are indicated. **C-D**. In silico footprinting using Tobias. Scatter plots of differential footprinting depth of transcription factor motifs in differentially accessible regions between WT and week 1 (C) and week 5 (D). Motifs with more footprint depth in WT are labelled in pink and in Taf4 mutants in light blue.

**Supplemental Figure 4.**
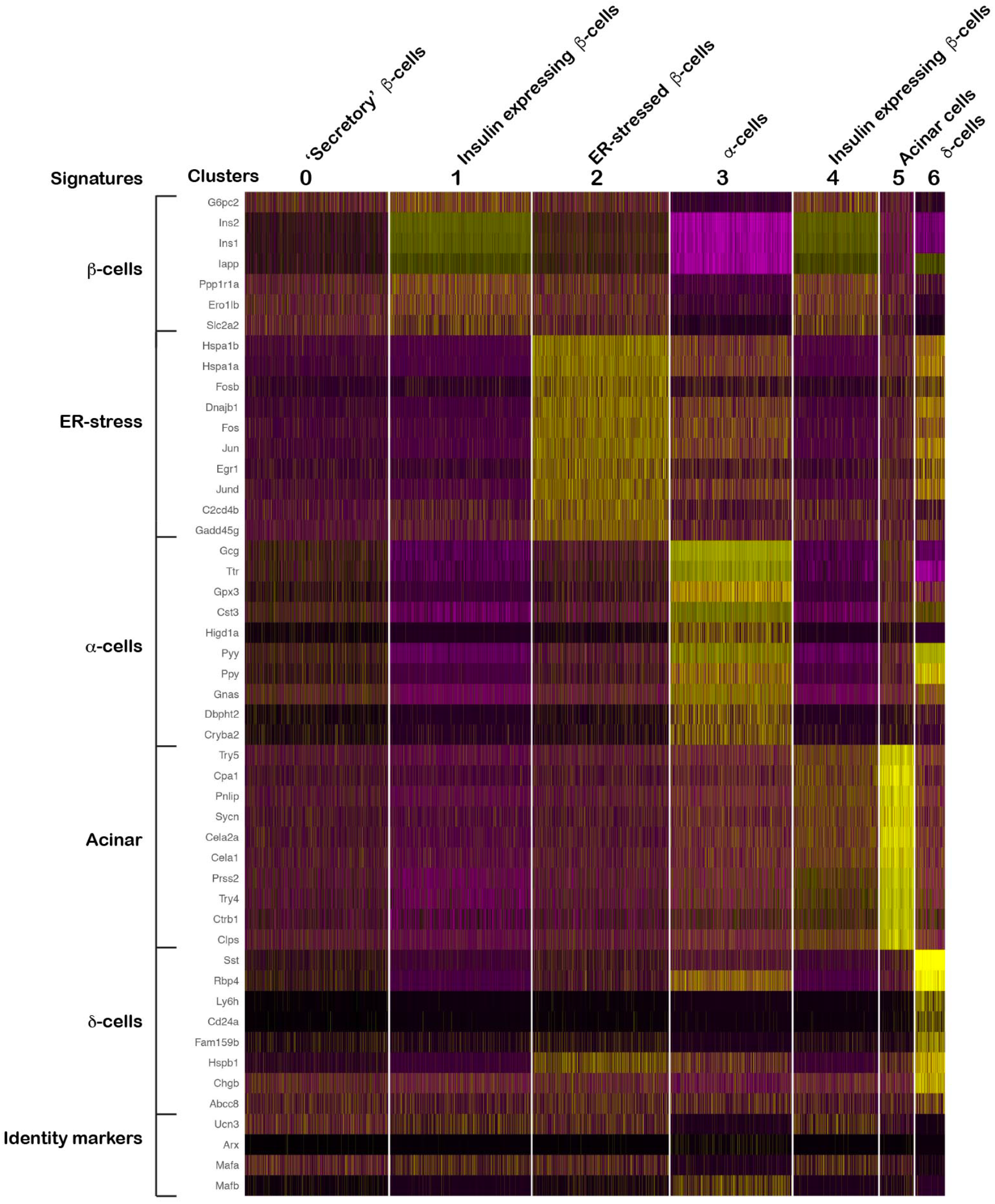
Heatmap of differentially expressed genes in the cell clusters from scRNA-seq of WT islets.

**Supplemental Figure 5.**
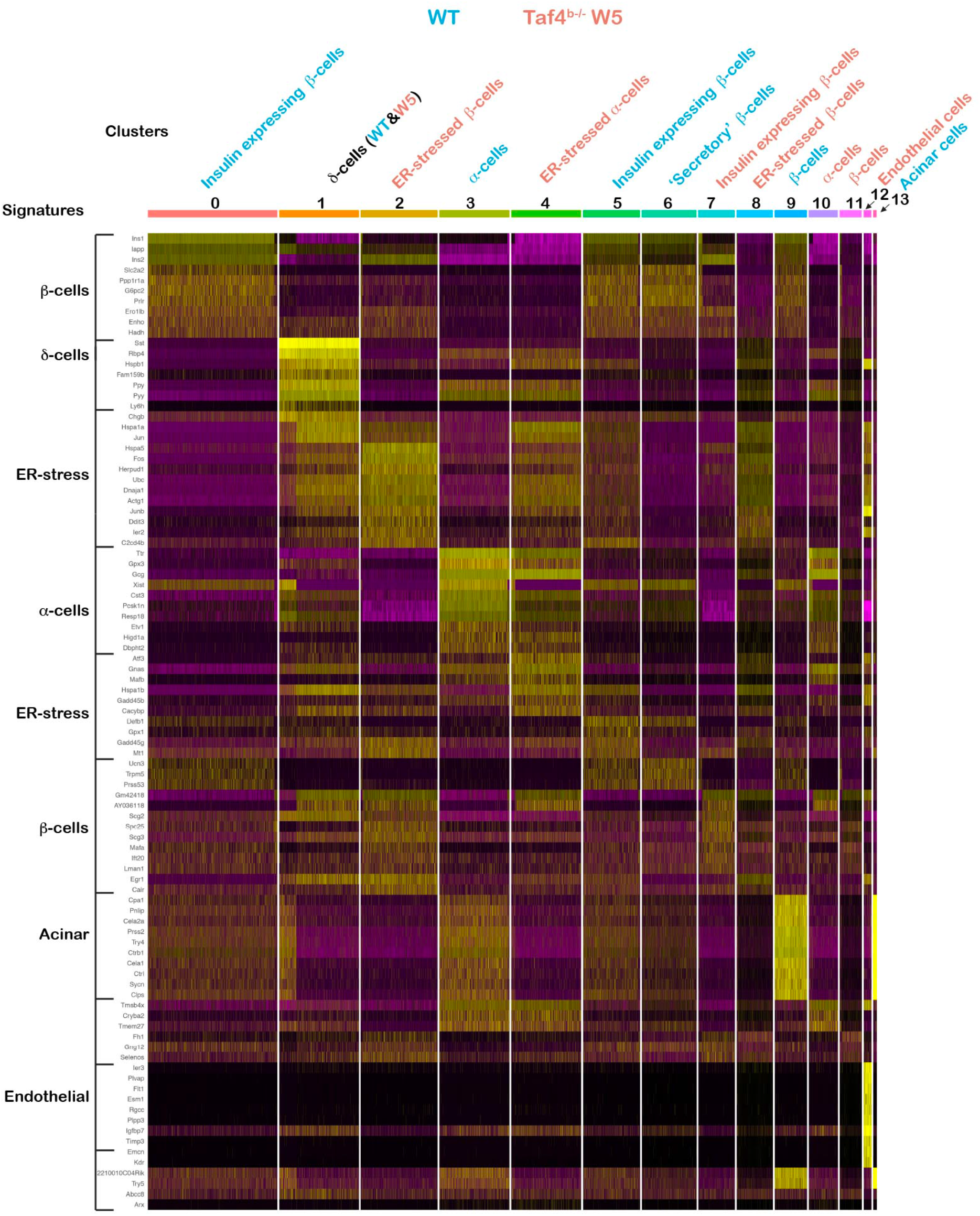
Heatmap of differentially expressed genes in the cell clusters from the WT/week 5 aggregate data.

**Supplemental Figure 6.**
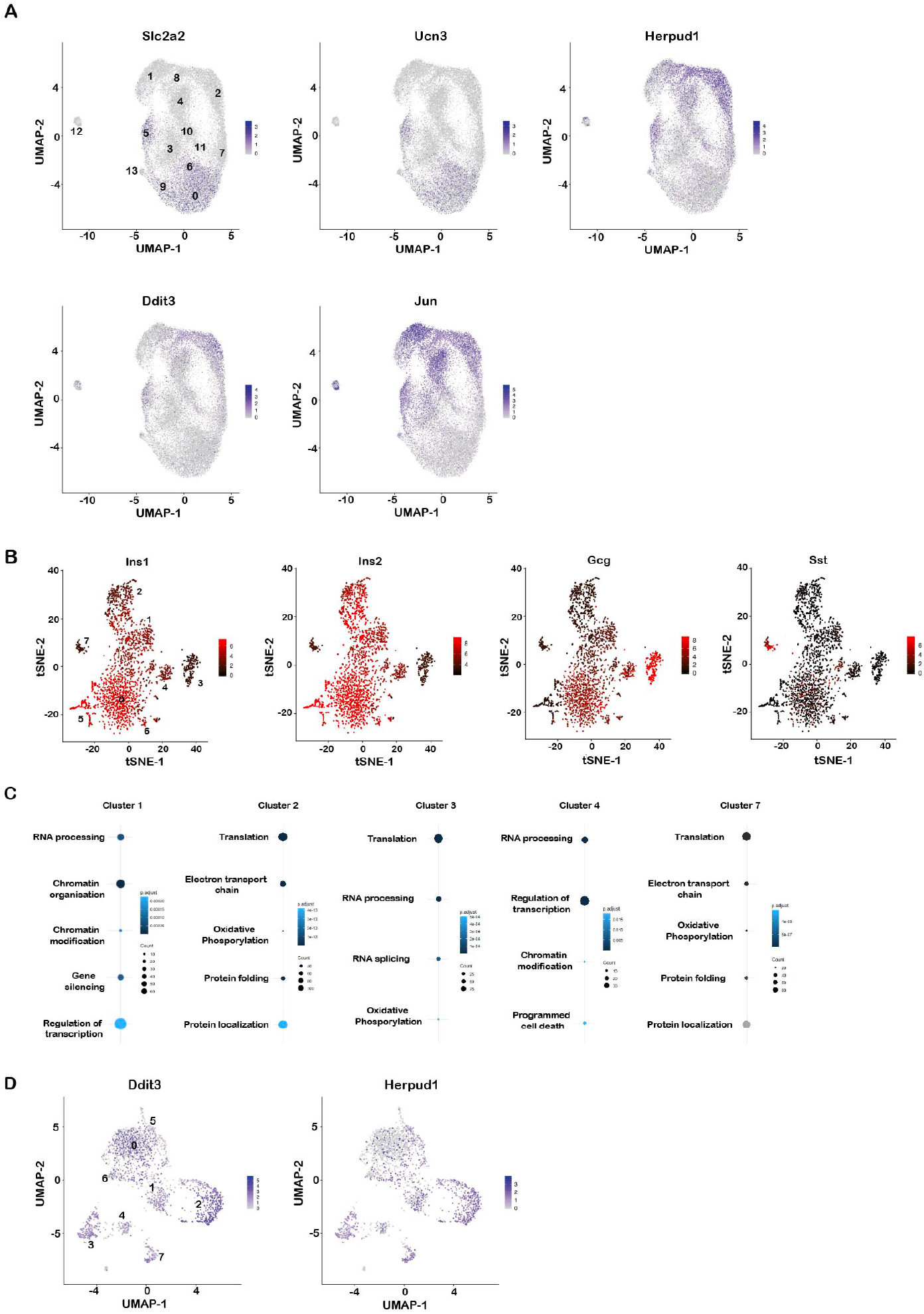
**A**. UMAP representations of the aggregate data illustrating the expression of the genes indicated in each panel. **B** tSNE representations of the week 1 cell populations illustrating the expression of the genes indicated in each panel. **C.** Bubble plots showing ontology (BP-FAT) of genes differentially expressed in the indicated clusters. **D**. UMAP representations week 1 cell populations illustrating the expression of the genes indicated in each panel.

**Supplemental Figure 7.**
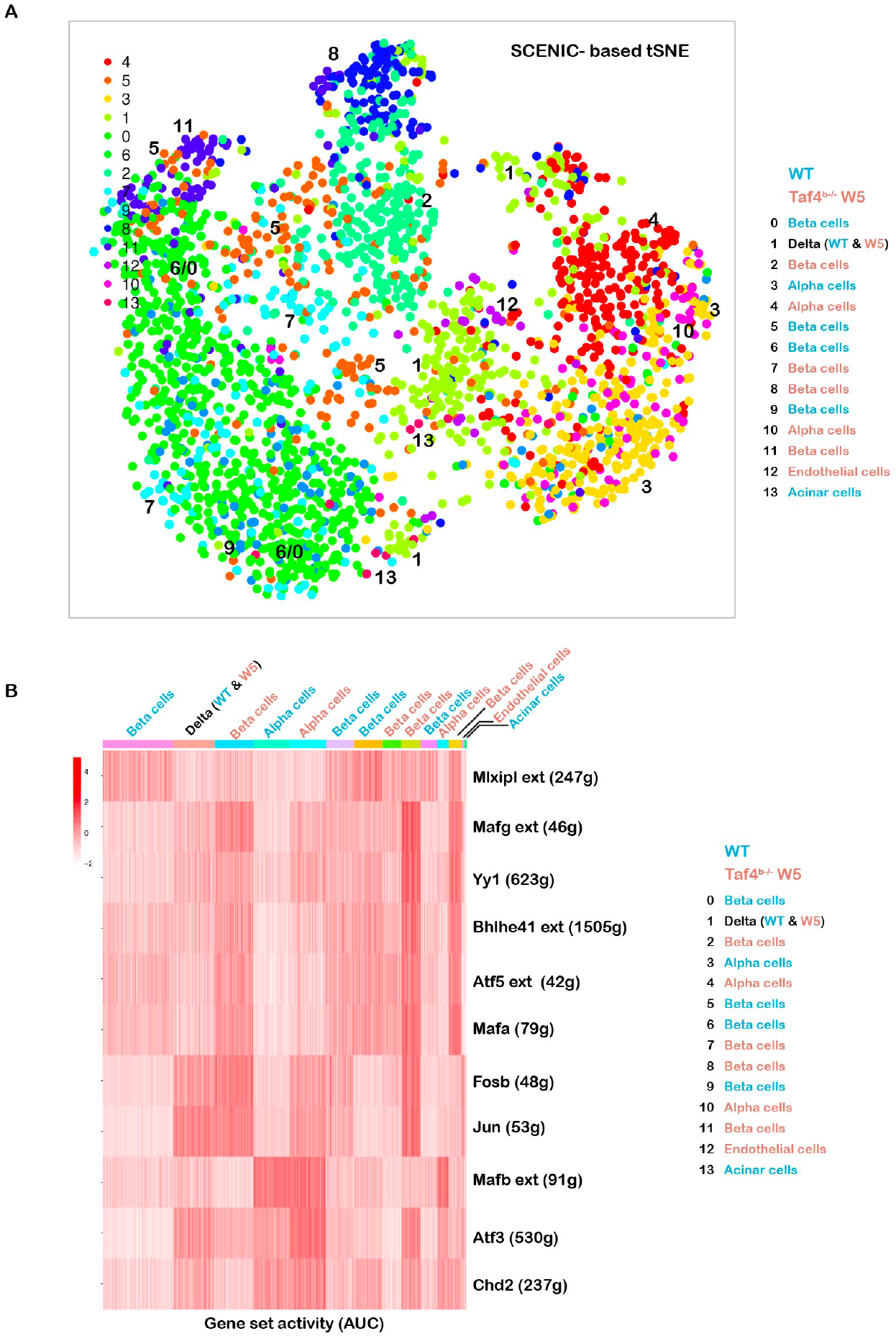
SCENIC analyses of WT and 5 week mutant islets populations. **A.** SCENIC based tSNE of 2500 cells from the WT/W5 aggregate. The identities of the cell populations are indicated. **B.** Heatmap representation of regulon activities in the different cell populations were quantified using AUCell.

**Supplemental Figure 8.**
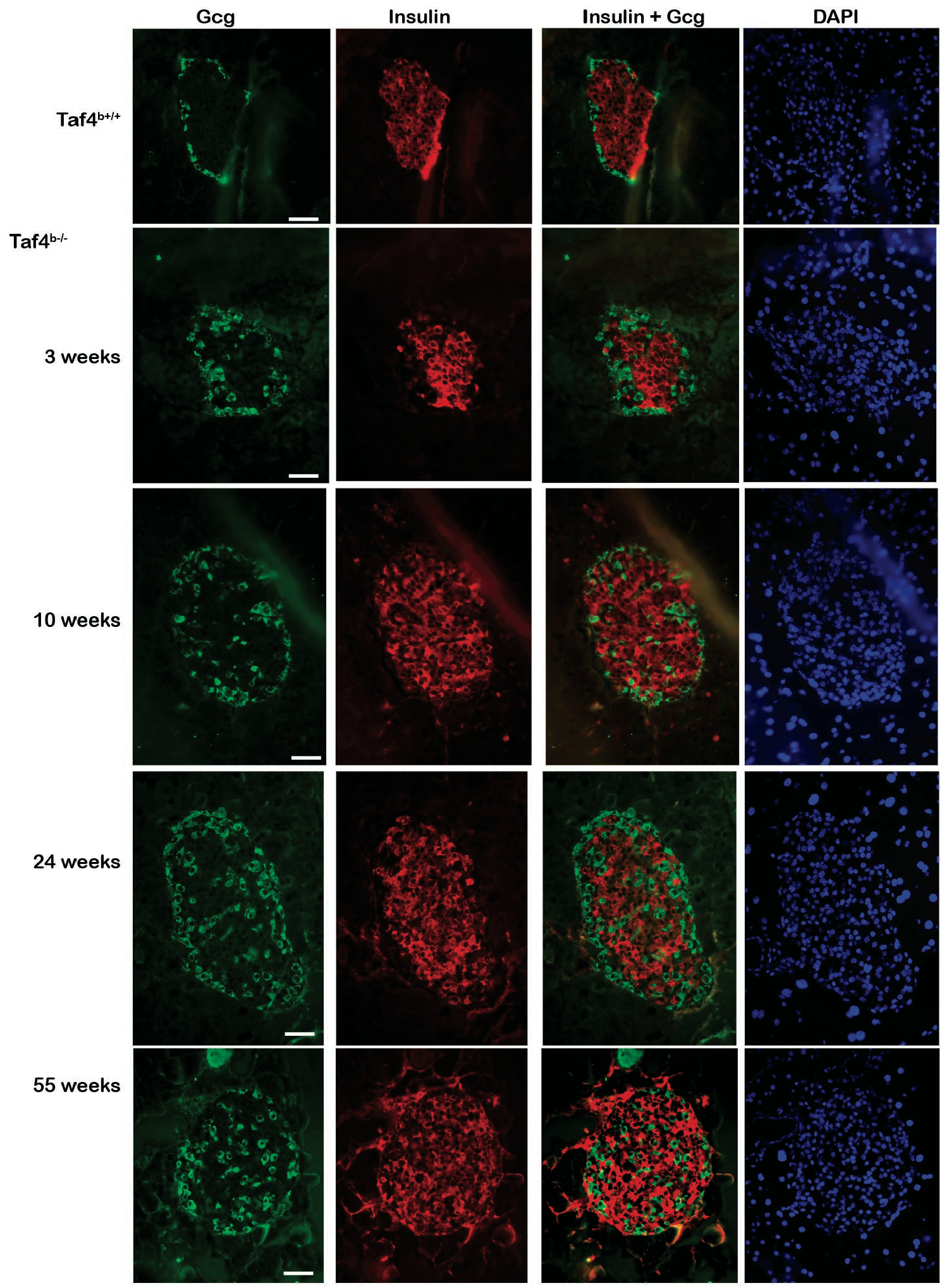
Immunostaining of isolated Langerhans islets from mice with the indicated genotypes for Insulin, Gcg or DAPI as indicated. The number of weeks after Tam injection are indicated. The Insulin-Gcg merge is shown to illustrate the mutually exclusive nature of the labelling. Scale bar = 100 μM

**Supplemental Dataset 1.** Summary of RNA-seq 1 3 and 5 weeks after Taf4 inactivation. Each spreadsheet shows the genes up (UR) and down-regulated (DR) along with the corresponding ontology analyses.

**Supplemental Dataset 2.** Summary of genes differentially regulated in the cell populations 1 week after Taf4 inactivation along with their ontologies.

## Notes

The authors declare no potential conflicts of interest.

### Competing Interest Statement

The authors have declared no competing interest.

### Summary of Updates

This version corrects an error in the description of Fig.2 and comprises a reworded Introduction and Discussion.

